# Why are fishers retaining manta and devil ray bycatch?

**DOI:** 10.64898/2026.09.01.748649

**Authors:** Mayuri Chopra, Divya Karnad, Gwilym Rowlands, Guy M. W. Stevens, Tejaswi Abhiram Nagam, T. Mohanraj, Daniel Fernando, Betty Laglbauer, Katrina J. Davis

## Abstract

Increasing fishing pressure, including from small-scale fisheries, has caused declines in more than one-third of all elasmobranch species. Tackling conservation issues in small-scale fisheries requires interdisciplinary approaches due to the coastal community’s interdependence on marine resources. To support inclusive policy change and fisher engagement, an understanding of the motivations driving fishers’ operational choices (*i.e.,* the retention of elasmobranch bycatch) is needed. We assess the motivational drivers behind bycatch retention of one of the slowest-growing and most vulnerable elasmobranch groups, manta and devil rays (collectively, mobulids), through a case study in India, their largest fishery in the world. We conducted a best-worst scaling survey in the fishery-intensive states of Tamil Nadu and Andhra Pradesh, which make significant contributions to mobulid landings on India’s east coast. Our results suggest that fishers exhibit varied motivations for retaining mobulid bycatch across states. Financial motivation to sell mobulids for additional revenue was the most important motivator for bycatch retention in both states. In Tamil Nadu, the top three motivators were all financially driven, whereas in Andhra Pradesh, the top three motivators included both financial and non-financial attributes, such as nutritional importance and storage optimisation. As the first socio-economic study of mobulid fisheries in India, we show that motivations underlying bycatch retention decisions vary geographically and may be influenced by cultural differences between states and the socio-economic characteristics of decision makers. Based on identified fisher motivations, we provide context-specific recommendations to align conservation strategies with the values fishers derive from the mobulid fishery and encourage participation in conservation. These include subsidies for net repair to encourage mobulid release; promotion of a minimum price measure for sustainably sourced alternative species; quality improvement of target species; and increased awareness of national and international regulatory obligations (e.g., CITES, CMS, IOTC).

## 3. Introduction

Overfishing has severe negative impacts on vulnerable marine populations globally, especially elasmobranchs (sharks, rays, and skates), due to their slow life histories (Stevens et al., 2000; Frisk et al., 2001; Frisk et al., 2005; Blaber et al., 2009; Davies et al., 2009). Increasing fishing pressure over the past decade has contributed to population declines in more than one-third of elasmobranch species, which are now facing elevated extinction risk (Davidson et al., 2016; Dulvy et al., 2021; Chopra et al. 2026b). The extinction risk of elasmobranchs is highest in coastal tropical waters, where small-scale fisheries (SSFs) are most prevalent, accounting for a significant proportion of threatened elasmobranch catches (Di Lorenzo et al., 2022). Overfishing in SSFs is increasingly recognised but remains challenging to manage due to capacity-limited enforcement systems, and limited catch regulation and reporting through management institutions (Cinti et al., 2014). The impact of SSFs on elasmobranchs through overfishing is also harder to manage due to coastal community dependencies on marine resources, making elasmobranch conservation in SSFs a complex socio-economic challenge (Béné and Friend, 2011; Weeratunge et al., 2013; Temple et al., 2024). Therefore, achieving sustainable elasmobranch fisheries requires an understanding not only of the biological drivers and environmental consequences of overfishing, but also the social and economic drivers shaping fisher behaviour (Temple et al., 2024).

Fisheries management improves with the inclusion of local communities because fishers’ decisions to target, retain, and trade certain species are embedded in socio-ecological systems and mediated by reciprocity and local institutions (Bajracharya et al., 2005; Sultana and Abeyasekera, 2008; Obradović et al., 2023). At the individual level, fisher decisions are influenced by a range of instrumental and non-instrumental motivations (Satumanatpan and Kanongdate, 2025). Consequently, ‘catch-ban’ policies for vulnerable species often fail to achieve effective fishery management outcomes because the problem remains unsolved, as fishers may still incidentally catch non-target species (Tolotti et al., 2015). Incentive-based conservation involving communities has gained global interest in positioning communities as stewards of conservation (Booth, 2021), however, designing of incentive-based programmes has fallen short of this rhetoric, largely due to missing nuances in evaluating the range of benefits required for success at the community level (Spiteri and Nepalz, 2006, Booth et al., 2019; Booth, 2026). Further, focusing purely on financial incentives may potentially have a negative impact by increasing catch risk and furthering unsustainable behaviours (Booth et al., 2025), in addition to concerns over their financial sustainability and scalability. Therefore, understanding the range of underlying motivations that lead to fishers’ decisions to target certain species is critical to designing effective incentive-based conservation interventions (Davis et al., 2017; Booth et al., 2020; Fordham et al., 2022).

While much previous research examining fishers’ motivations has focused on using methods such as interviews (Seidu et al., 2022; Young et al., 2016), these qualitative methods have drawbacks due to biases that increase outcome uncertainty for conservation (Small and Cook, 2023). Methods that include quantitative, structured, and randomised designs can overcome some of the biases of traditional interviews (Jervis and Drake, 2014). Best-worst scaling (BWS), a type of discrete choice experiment, is one such method that provides significant improvements over previous methods by providing statistically robust results with relatively smaller sample sizes and lower survey fatigue (Potoglou et al., 2011). Best-worst scaling has been widely used to understand motivations and perceptions in agriculture, transport, and healthcare (Mühlbacher et al., 2016; Teffo et al., 2019; Caputo and Lusk, 2020) and is increasingly being adopted in wildlife conservation to address complex socio-economic challenges (Davis et al., 2017; Tyner and Boyer, 2020; Yin et al., 2023; Schuster et al., 2024). The objective of our study is to identify fishers’ motivations for bycatch retention.

Manta and devil rays (collectively, mobulids) are an ideal study system for understanding choice behaviour in bycatch retention due to the urgent conservation need to address their global population declines from overfishing (Lawson et al., 2017; Jabado et al., 2025a, 2025b, 2025c), and their continued prevalence as secondary catch in SSFs worldwide (Croll et al., 2016; Laglbauer et al. 2026). The Indian Ocean region is a particularly important site to address fisher motivations for mobulid catch due to high population declines in Indian Ocean fisheries (Laglbauer et al., 2026), such as in Myanmar (Segura-García et al., 2024), Sri Lanka (Fernando and Stewart, 2021), Bangladesh (Haque et al., 2021), Indonesia (Lewis et al., 2015), and India (Chopra et al., 2026a). We used a BWS survey of fishers landing manta and devil rays in the study area to assess their motivations for retaining mobulid bycatch. In addition to understanding fishers’ perspectives and the importance of motivations for mobulid bycatch retention, we also examined how the socio-economic characteristics of fishers influence their choice behaviour (Novak Colwell and Axelrod, 2017). Our work informs conservation and sustainable management of manta and devil rays by providing broader insights into designing community-based behaviour change interventions and incentive-based conservation policies for threatened species.

## 4. Methods

### 4.1 Study site

Our case study system was the mobulid fishery in India. India ranks among the top five conservation priority nations for mobulids and hosts the world’s largest mobulid fishery (Palacios et al., 2025; Laglbauer et al., 2026), making it a critical case study for understanding driving motivators of bycatch retention. India also supports a coastal population of approximately 250 million people (UNISDR and UNDP, 2012) including a marine fisher population of approximately four million individuals, with 1.5 million actively engaged fishers and 2.5 million engaged in fishing-associated sectors (Ministry of Agriculture and CMFRI, 2012). Notably, about 61% of this population is economically marginalised and falls below the poverty line (Kumar and Shivani, 2014). Socio-economically vulnerable coastal communities in India rely heavily on marine resources, demonstrating the interdisciplinary nature of challenges in elasmobranch conservation (Umamaheshwari et al., 2021). Manta and devil rays are one such group of elasmobranchs facing this interdisciplinary conservation challenge due to being caught as opportunistic catch (referred to as bycatch in this study) in non-selective fishing gear (Croll et al., 2016) and forming a portion of income or sustenance for communities in India (Sultana et al., 2014; Karnad et al., 2020; Kizhakudan et al., 2024).

We surveyed the southeastern Indian states of Tamil Nadu and Andhra Pradesh, as they contribute over 86% to the east coast mobulid landings (CMFRI, 2015; Nair et al., 2015; Figure 1). Additionally, both states have a high proportion of the active fisher population relying on shark and ray catch for income and subsistence needs (CMFRI, 2015; Jabado et al., 2024, CMFRI, 2015). To identify fisher motivations, we surveyed fishers landing mobulids at prominent fish landing centres contributing to high mobulid bycatch (Nair et al., 2015; Chopra et. al, 2026a) at six data collection sites: three in Tamil Nadu (Threspuram Fishing Village [(8.8165° N, 78.1625° E]), Tharuvaikulam Fishing Harbour [(8.8922° N, 78.1707° E]), and Tuticorin Fishing Harbour [(8.7945° N, 78.1584° E]), and three in Andhra Pradesh (Visakhapatnam Fishing Harbour [(17.6868° N, 83.2185° E]), Pudimadaka Fishing Village [(17.4927° N, 83.0028° E]), and Kakinada Fishing Harbour [(16.9891° N, 82.2475° E]) (Figure 1). Fishers landing mobulids in our study sites engaged in target fisheries dominated by tuna, other pelagic fish including seer fish (*Scombridae*), murrel fish (*Channidae*), sailfish (*Istiophoridae*), and other sharks and rays (Chopra et al., 2026a).

**Figure 1.**
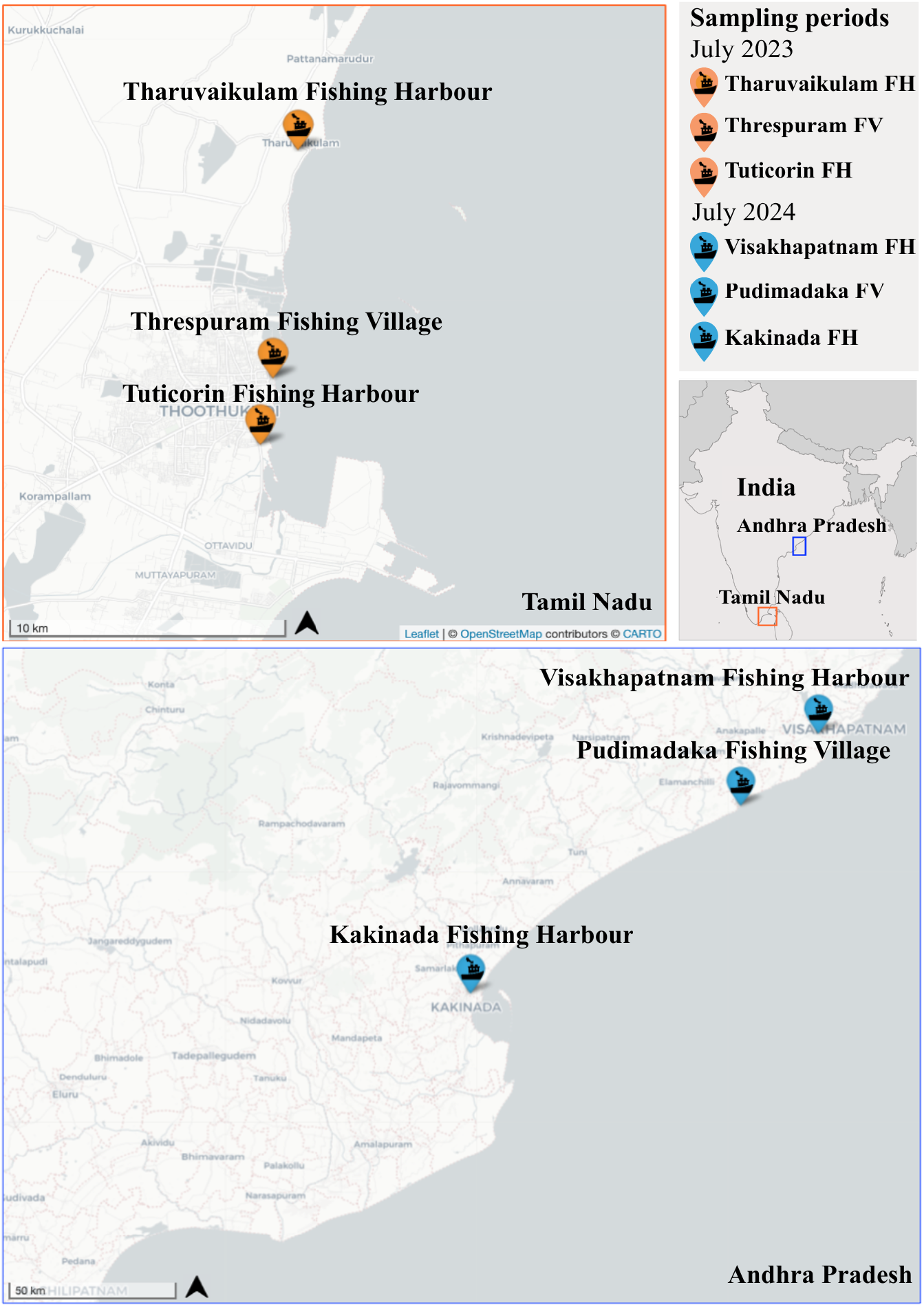
Map of study sites along the east coast of India. The study sites are in the states of Tamil Nadu (Tharuvaikulam Fishing Harbour, Threspuram Fishing Village and Tuticorin Fishing Harbour) and Andhra Pradesh (Visakhapatnam Fishing Harbour, Pudimadaka Fishing Village and Kakinada Fishing Harbour). Surveys were conducted with fishers at these locations to identify the motivations for bycatch retention of manta and devil rays (FH: fishing harbour; FV: fishing village).

### 4.2 Data collection

#### 4.2.1 Survey development and administration

We assessed key operational behaviours driving elasmobranch bycatch retention in fisheries, and thereby impacting fisheries sustainability. Our survey had three sections. In section one, we recorded respondents’ fishing characteristics including role in fishing, boat size, age, years of fishing experience, duration of fishing, and fishing distance from the coast. In section two, we presented a BWS experiment to assess respondents’ most and least important motivations for retaining mobulid bycatch (Table 1). In section three, we collected socio-economic data such as monthly income, whether respondents were the main earner in the household, and formal education level. The survey was conducted face-to-face over 40 survey days in July 2023 and July 2024. Sampling was prioritised in July, during the peak tuna fishing season (Kumar, 2017), as this is when the majority of mobulids are landed. Fishers were approached opportunistically at fish landing centres and asked if they landed manta and devil rays, and only consenting individuals who responded with ‘yes’ were selected as survey respondents (ethics under University of Oxford: CUREC REF R85742/RE001). We conducted surveys using a representative sample of the active fisher population across selected sites. In Tamil Nadu, the surveyed locations included Threspuram Fishing Village (with 3,643 active fishers), Tharuvaikulam Fishing Harbour (with 1,559 active fishers), and Tuticorin Fishing Harbour (with 2,881 active fishers) (CMFRI-DoF, 2020a), with respondents constituting a representative proportion of approximately 1.21% of the state’s active fisher population. In Andhra Pradesh, surveys were conducted in Visakhapatnam Fishing Harbour (with 3,931 active fishers), Pudimadaka Fishing Village (with 202 active fishers), and Kakinada Fishing Harbour (with 325 active fishers) (CMFRI-DoF, 2020b), where respondents constituted a representative proportion of approximately 1.9% of the active fisher population.

**Table 1.**
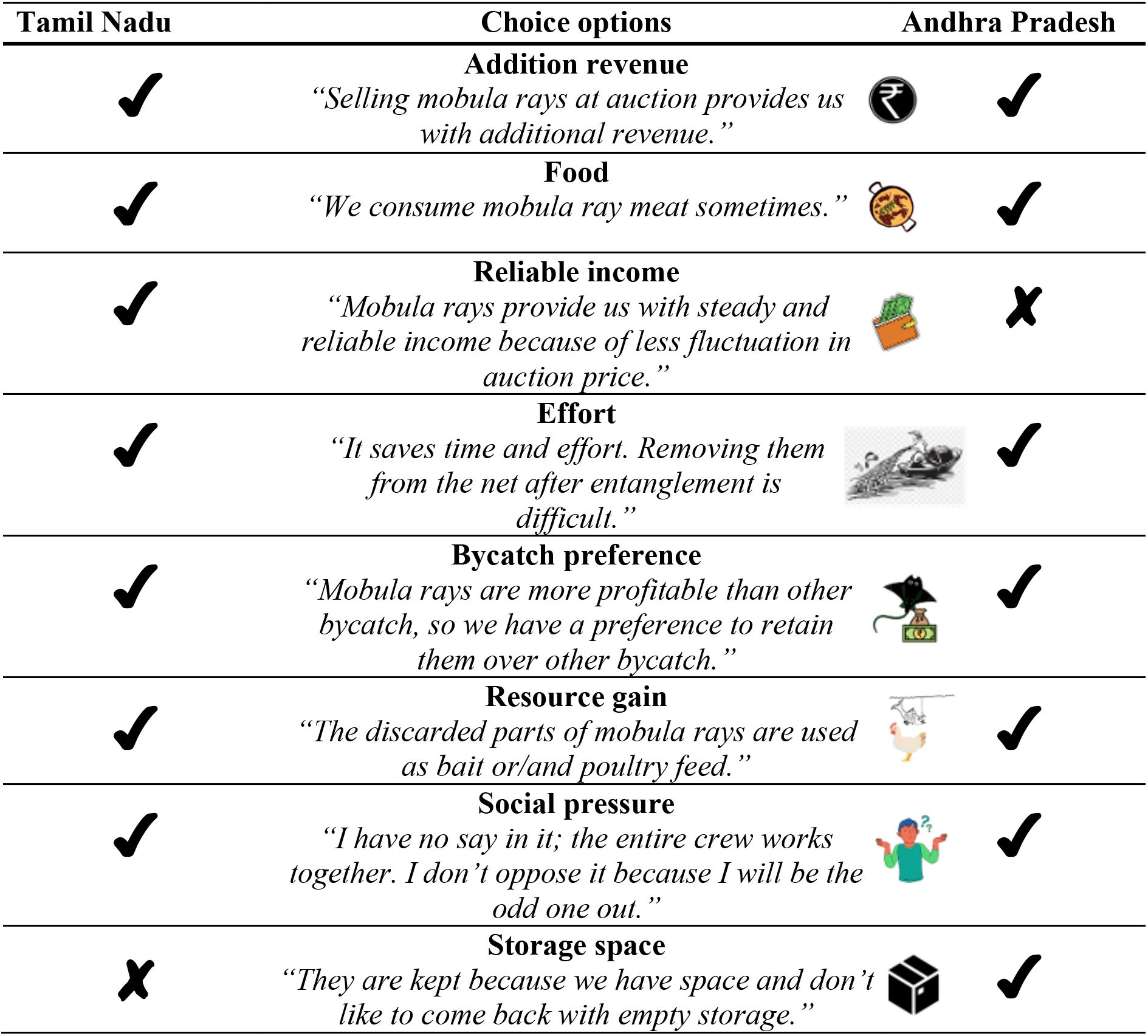
Options used to identify motivations for mobulid bycatch retention tested in a best-worst scaling surveys with fishers in Tamil Nadu and Andhra Pradesh. The tick indicates the options used in the surveys conducted in the location and crosses indicate the options eliminated in the state during pilot surveys.

#### 4.2.2 Best-worst scaling

To assess fisher motivations driving mobulid bycatch retention, we used BWS, a type of discrete choice experiment (Finn and Louviere, 1992), in which we presented respondents with choice sets from which they chose the ‘best’ and ‘worst’ options (example in Figure 2). Parametric analyses of choice data using best-fit models can predict how often an option will be chosen over another based on Random Utility Theory (Flynn and Marley, 2014). We use BWS because it offers two methodological advantages that are particularly relevant to our study. First, BWS imposes a lower cognitive burden than traditional ranking tasks (Potoglou et al., 2011), an important consideration given that most fisher respondents have not received formal education (CMFRI-DoF, 2020a; 2020b) and were surveyed in busy port environments. Second, BWS can generate statistically robust results with relatively small sample sizes due to the orthogonality inherent in the balanced block designs used in BWS surveys (Flynn et al., 2008). This was a key benefit in our study, as in-person data collection constrained the number of individuals that could be reached (e.g., compared to panel data sets). We employed a Case I BWS, where we assumed that the chosen ‘best’ and ‘worst’ choices represented the maximum difference in appeal for respondents (Louviere and Flynn, 2010).

**Figure 2.**
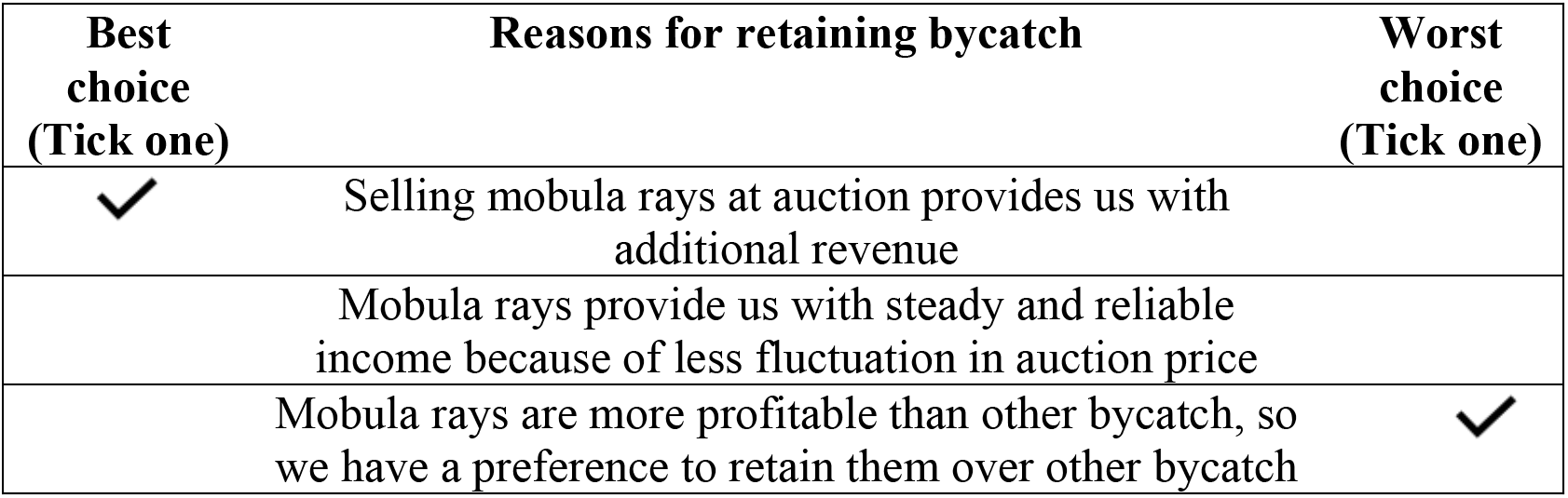
An example of a best-worst scaling choice set from the conducted fisher surveys. The fisher surveys were conducted in Tamil Nadu and Andhra Pradesh to identify fisher motivations behind mobulid bycatch retention. Note: This is an English original version which was translated in Tamil or Telugu languages.

To design the BWS experiment, we developed a list of eleven potential motivators driving shark and ray bycatch retention in small-scale fisheries (SSFs) (Supplementary Information 1). To reduce respondent fatigue, we engaged seven experts from the field of shark and ray conservation to rank the list of options, following which the two lowest ranking motivations were removed. Additionally, we reduced the number of potential motivators by testing options in a focus group discussion (5–7 participants) and conducting five pilot surveys across both states. Results from this discussion and testing left us with a final list of seven state-specific options (Table 1; Supplementary Information 1). We used the shortlisted seven motivations in a balanced incomplete block design (BIBD) to generate seven choice sets, each containing three choice options, which co-occurred once with every option (Louviere et al., 2015). To avoid order bias (Campbell and Erdem, 2015), we ensured random ordering of choice options in each set, with no option consistently appearing first, middle, or last.

### 4.3 Data analysis

To assess motivations for retaining mobulid bycatch, we first used the best-worst scores to rank preferences in Tamil Nadu and Andhra Pradesh using non-parametric analyses (Supplementary Information 2). Best worst scores were calculated as the scores of an item (*i*) based on the number of times respondent (*n*) selected it as the best (B*_in_*) or the worst (W*_in_*) choice relative to opportunity (Aizaki and Fogarty, 2023). To model choice responses using parametric analyses, we used the Maxdiff model, assuming that fisher respondents select item *i* and *j* from a choice set where *i* is selected as the best motivator and *j* is selected as the worst motivator, and the difference in utility between *i* and *j* is the maximum among all utility differences (Hensher et al., 2015). We defined the probability of the fisher respondent choosing item *i* as best and *j* as worst using the conditional logit framework (Aizaki and Fogarty, 2023), as:

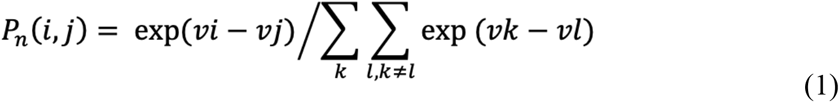

Where *P_n_* is the probability of selecting an item, *v_i_* is the systematic component of the utility of item *i*, defined by the product of a coefficient to be estimated and a dummy variable (with a value of 1 when included in the choice set and 0 when not included) (Aizaki and Fogarty, 2023). To comparatively assess all bycatch retention motivators, the reference (model base case) in the conditional logit model was set as the option ‘Additional revenue: *Selling mobula rays at auction provides us with additional revenue*’, due to it being the highest-ranking motivator across both states. To assess the impact of respondent socio-demographics on bycatch retention choice behaviour, we added interaction terms to the base conditional logit model. We nested the base conditional logit model within the full model and retained interaction terms that significantly improved model fit (log-likelihood ratio test, p < 0.05) using backward elimination.

As models of best choices often differ from models of worst choices due to greater consistency in negative evaluations than in positive ones, we expected differences between the best and worst choice behaviour (Czapiński and Lewicka, 1979; Flynn and Marley, 2014; Rigby et al., 2015; Davis et al., 2019). To assess whether our best and worst choice responses could be modelled together, we tested consistency in choice rankings using a log-likelihood ratio test. Finally, we interpreted probabilities for bycatch retention motivators using the log-odds ratios of conditional logit model estimates (Lipovetsky and Conklin, 2014). We analysed all data using RStudio Version 2024.12.0+467 (R Core team, 2024) and used the packages support.bws (Aizaki and Fogarty, 2023), crossdes (Sailer, 2013), dfidx (Croissant, 2020a), survival (Therneau and Grambsch, 2000) and mlogit (Croissant, 2020b).

## 3. Results

### 5.1 Socio-economic characteristics of mobulid fishers

We surveyed a total of 211 fishers across Tamil Nadu (n = 114) and Andhra Pradesh (n = 97). Missingness in responses prevented us from using the entire dataset in conditional logit modelling and we used 86% of the surveyed data (n = 181) in our statistical analyses (Tamil Nadu, n = 98 and Andhra Pradesh, n = 83; Table 2). All respondents were male, as only males are involved in active offshore fishing in both states (CMFRI-DoF, 2020a; 2020b). The majority of respondents in both states were boat workers (40% in Tamil Nadu and 42% in Andhra Pradesh), aged between 26 to 55, and involved in fishing trips lasting two to ten days (Tamil Nadu: 57%; Andhra Pradesh: 35%) (Table 2). Most respondents (Tamil Nadu: 99%; Andhra Pradesh: 92%) were the main earners of their family (Table 2). Over 70% of the fishers in both states reported a monthly income of under INR 30,000 (∼£250; GBP conversion rate = INR 121) (Table 2). Andhra Pradesh fishers utilised the coastal zones under 25 nautical miles, unlike fishers from Tamil Nadu, and a proportion of fishers from both states fished beyond the Exclusive Economic Zone (Tamil Nadu fishers: 21%; Andhra Pradesh fishers: 28%). A higher proportion of fishers in Andhra Pradesh had no formal schooling compared to Tamil Nadu (62% and 24%, respectively). In Tamil Nadu, none of the respondents were educated beyond the 12th grade, whereas in Andhra Pradesh, about 15% had completed schooling beyond the 10th grade and 4% were graduates engaged in fishing practices (Table 2).

**Table 2.**
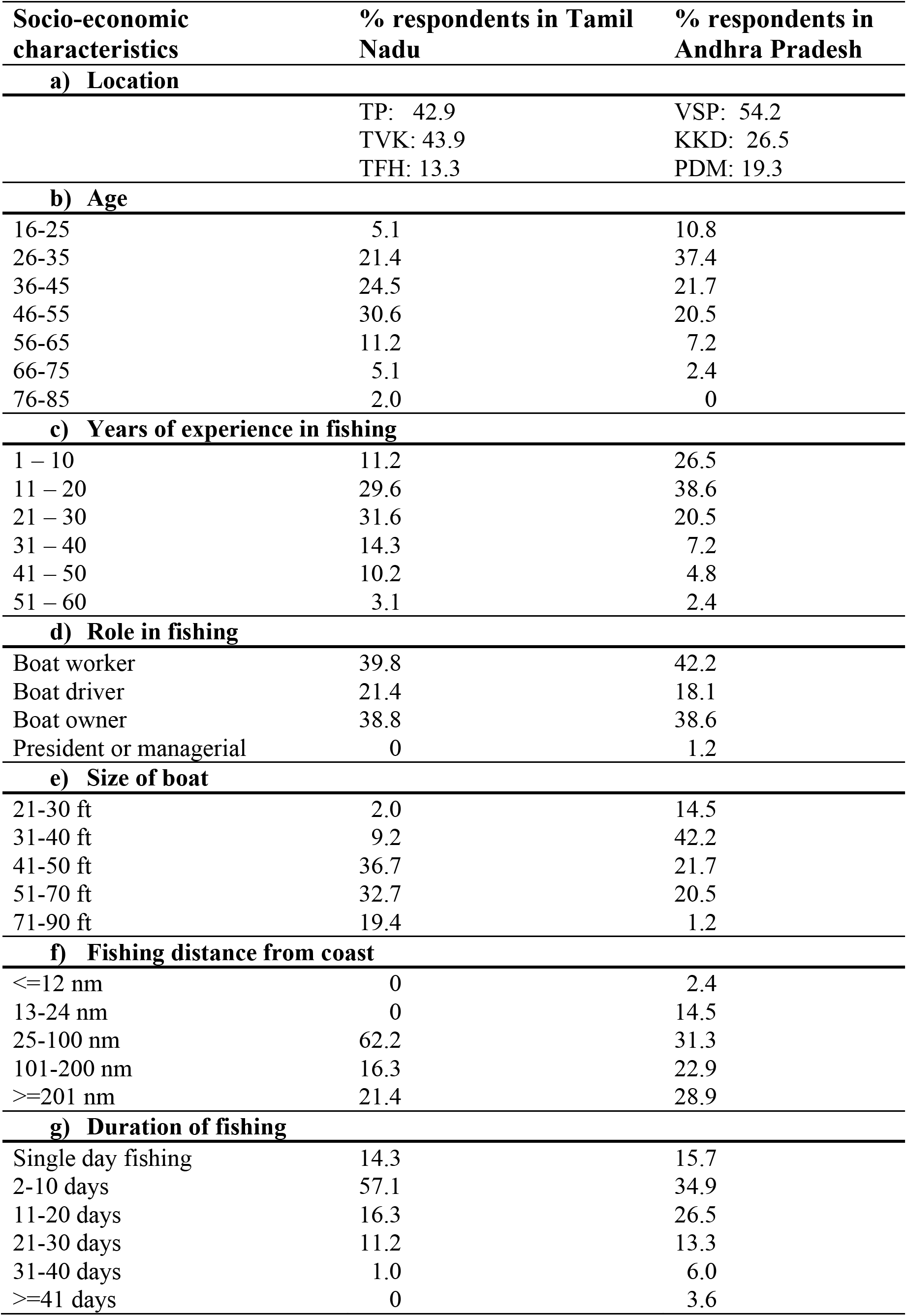
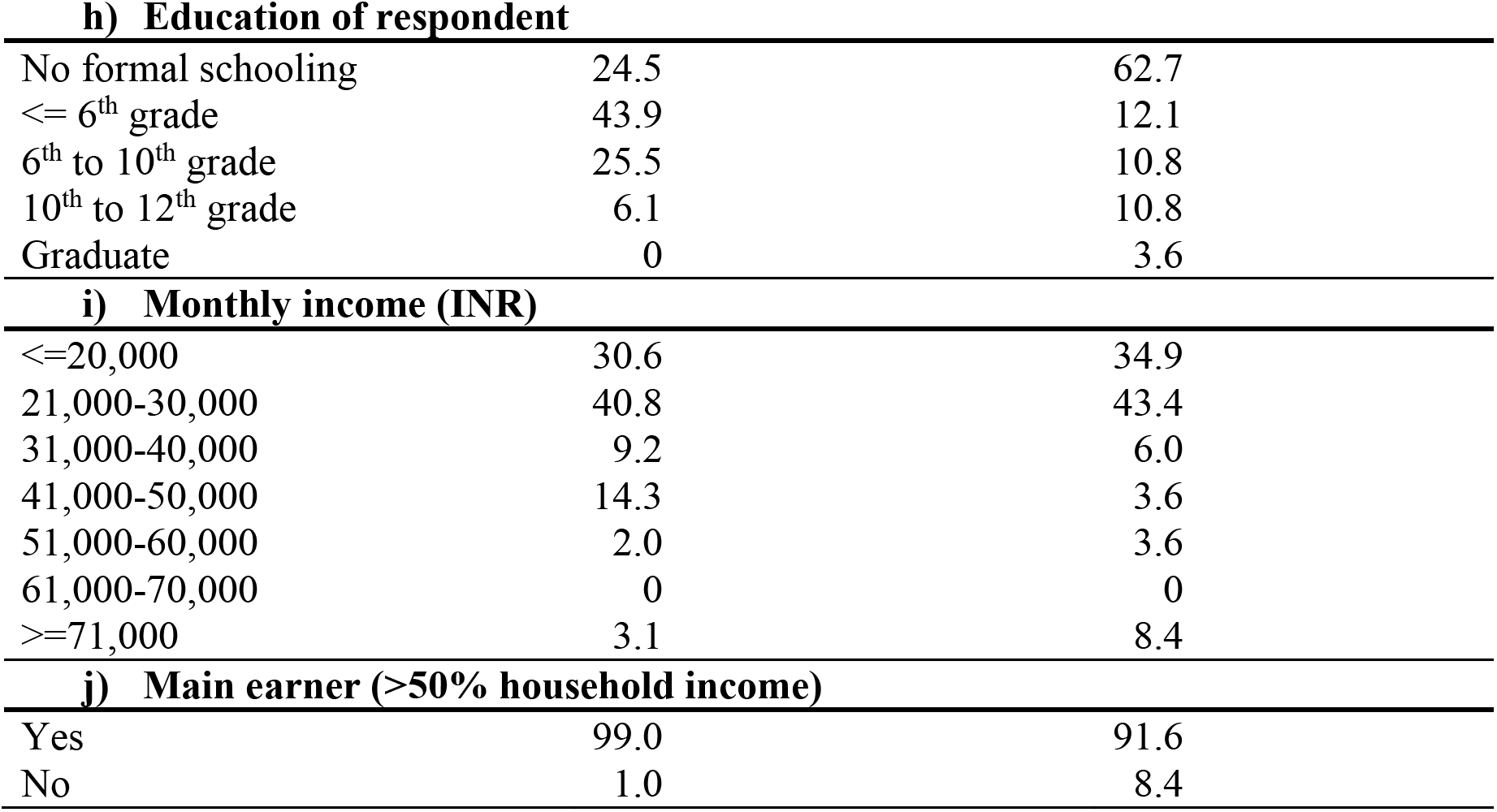
Socio-economic characteristics of the fisher respondents for the best-worst scaling survey conducted at study sites in India. Study states include Tamil Nadu (n = 98) and Andhra Pradesh (n = 83) from Threspuram Fishing Village (TP), Tharuvaikulam Fishing Harbour (TVK), Tuticorin Fishing Harbour (TFH), Visakhapatnam Fishing Harbour (VSP), Kakinada Fishing Harbour (KKD), and Pudimadaka Fishing Village (PDM).

| **Socio-economic characteristics** | **% respondents in Tamil Nadu** | **% respondents in Andhra Pradesh** |
| --- | --- | --- |
| **a) Location** |  |  |
|  | TP: 42.9 | VSP: 54.2 |
|  | TVK: 43.9 | KKD: 26.5 |
|  | TFH: 13.3 | PDM: 19.3 |
| **b) Age** |  |  |
| 16-25 | 5.1 | 10.8 |
| 26-35 | 21.4 | 37.4 |
| 36-45 | 24.5 | 21.7 |
| 46-55 | 30.6 | 20.5 |
| 56-65 | 11.2 | 7.2 |
| 66-75 | 5.1 | 2.4 |
| 76-85 | 2.0 | 0 |
| **c) Years of experience in fishing** |  |  |
| 1 – 10 | 11.2 | 26.5 |
| 11 – 20 | 29.6 | 38.6 |
| 21 – 30 | 31.6 | 20.5 |
| 31 – 40 | 14.3 | 7.2 |
| 41 – 50 | 10.2 | 4.8 |
| 51 – 60 | 3.1 | 2.4 |
| **d) Role in fishing** |  |  |
| Boat worker | 39.8 | 42.2 |
| Boat driver | 21.4 | 18.1 |
| Boat owner | 38.8 | 38.6 |
| President or managerial | 0 | 1.2 |
| **e) Size of boat** |  |  |
| 21-30 ft | 2.0 | 14.5 |
| 31-40 ft | 9.2 | 42.2 |
| 41-50 ft | 36.7 | 21.7 |
| 51-70 ft | 32.7 | 20.5 |
| 71-90 ft | 19.4 | 1.2 |
| **f) Fishing distance from coast** |  |  |
| <=12 nm | 0 | 2.4 |
| 13-24 nm | 0 | 14.5 |
| 25-100 nm | 62.2 | 31.3 |
| 101-200 nm | 16.3 | 22.9 |
| >=201 nm | 21.4 | 28.9 |
| **g) Duration of fishing** |  |  |
| Single day fishing | 14.3 | 15.7 |
| 2-10 days | 57.1 | 34.9 |
| 11-20 days | 16.3 | 26.5 |
| 21-30 days | 11.2 | 13.3 |
| 31-40 days | 1.0 | 6.0 |
| >=41 days | 0 | 3.6 |
| **h) Education of respondent** |  |  |
| No formal schooling | 24.5 | 62.7 |
| <= 6<sup>th</sup> grade | 43.9 | 12.1 |
| 6<sup>th</sup> to 10<sup>th</sup> grade | 25.5 | 10.8 |
| 10<sup>th</sup> to 12<sup>th</sup> grade | 6.1 | 10.8 |
| Graduate | 0 | 3.6 |
| **i) Monthly income (INR)** |  |  |
| <=20,000 | 30.6 | 34.9 |
| 21,000-30,000 | 40.8 | 43.4 |
| 31,000-40,000 | 9.2 | 6.0 |
| 41,000-50,000 | 14.3 | 3.6 |
| 51,000-60,000 | 2.0 | 3.6 |
| 61,000-70,000 | 0 | 0 |
| >=71,000 | 3.1 | 8.4 |
| **j) Main earner (>50% household income)** |  |  |
| Yes | 99.0 | 91.6 |
| No | 1.0 | 8.4 |

### 5.2 Fisher perception of the mobulid fishery

We recorded fisher perceptions for 111 respondents in Tamil Nadu (97% of respondents) and 37 respondents in Andhra Pradesh (38% of respondents). Fisher perceptions of the mobulid fishery varied between the two states (Figure 3). In Tamil Nadu, the majority of the fishers (>10 fishers for each word description) perceived a mobulid sighting as ‘income’, and were encouraged to ‘lift’ (referring to pulling up mobulids entangled in the net onto the vessel). In Andhra Pradesh, the fisher perceptions were more diverse, indicating a broader range of associations beyond sole economic value. The majority of fishers in Andhra Pradesh perceived mobulid sightings as ‘beautiful’, ‘big’, ‘powerful’, ‘fear’, ‘danger’, ‘cheap’, ‘catch’, ‘nothing’ (>1 fisher for each word description).

**Figure 3.**
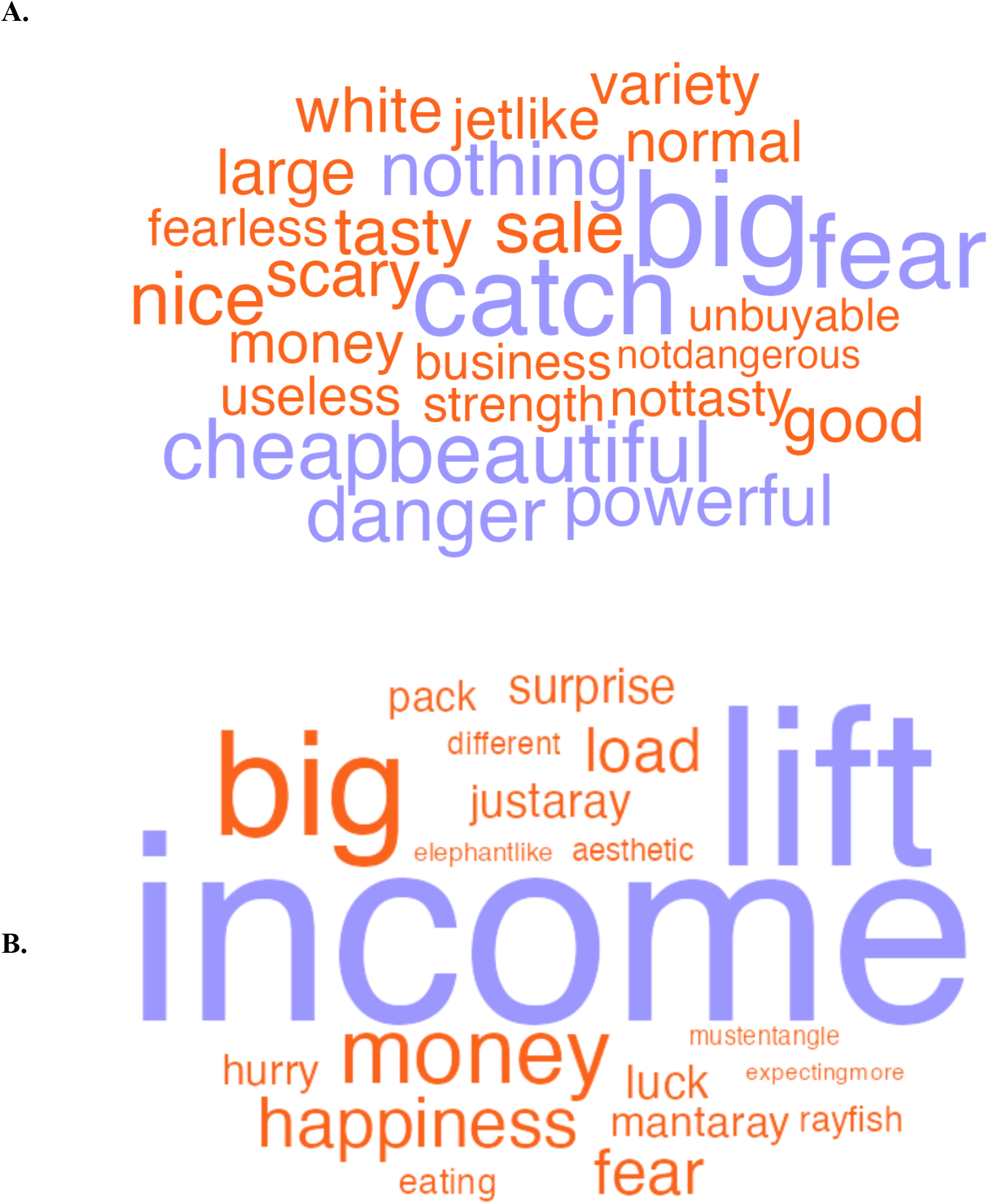
A world cloud representing perceptions of respondent fishers on manta and devil rays in the east coast of India. **A.** Andhra Pradesh responses from 37 fishers (38% of surveyed respondents). Purple words in A. show words recorded more than once (maximum thrice) and orange words are those recorded just once. **B.** Tamil Nadu responses from 111 fishers (97% of surveyed respondents). Purple words in B. are words with frequency over ten times, and orange words are those recorded under ten times.

### 5.3 Best-worst scaling analyses

The log-likelihood ratio testing consistency between best and worst responses showed a significant difference between the (best-worst) separate and the (best-worst) combined models (p < 0.001), indicating that the best and the worst responses should be modelled separately. Therefore, we only present the results of the best responses here, consistent with previous literature (Schuster et al., 2024). The ranking of the top three motivators remained unchanged between the non-parametric analysis (best-worst score estimates) and the parametric best-response analyses for both states (Table 3 and Table 4).

**Table 3.**
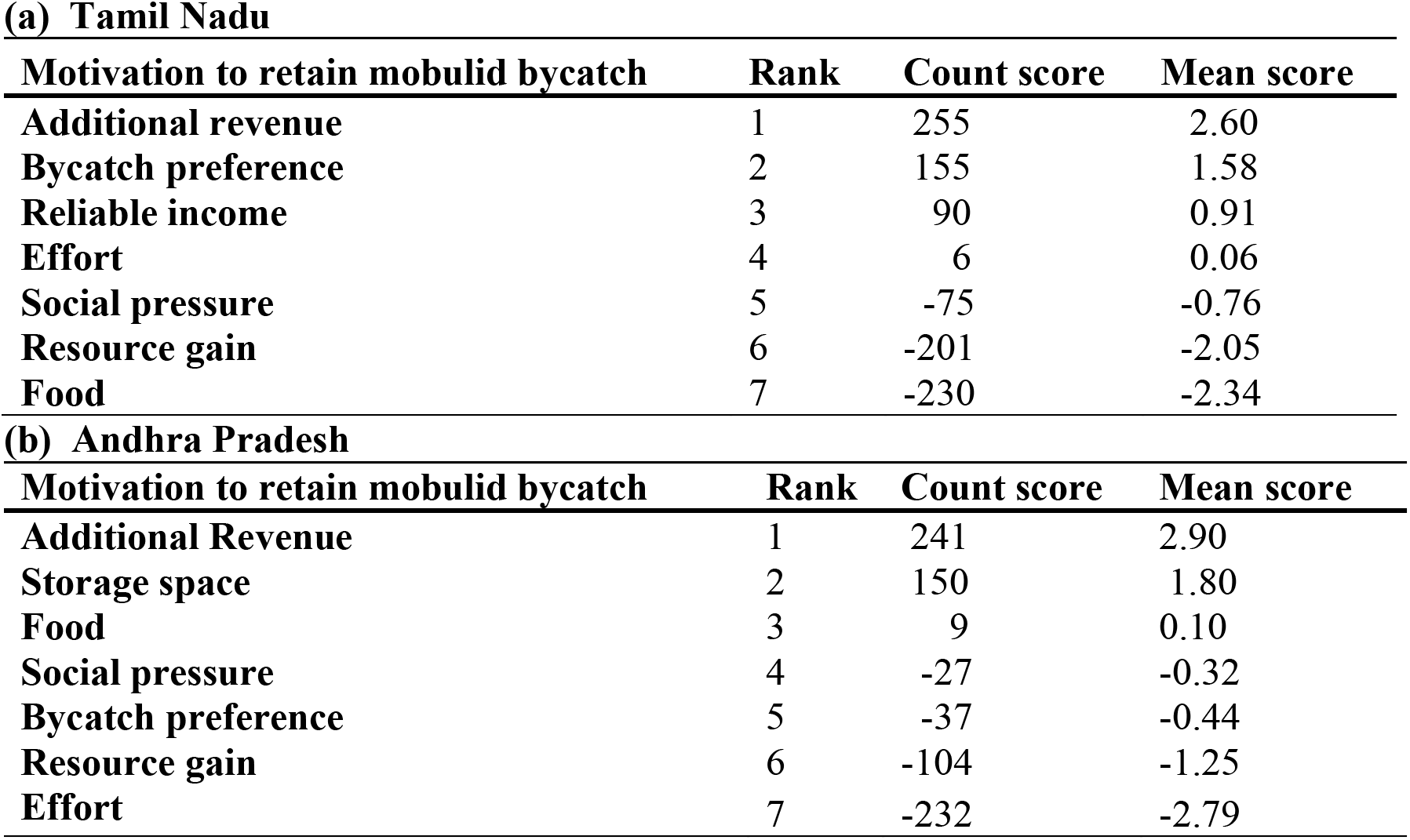
Best-worst scores calculated from fisher choice responses to the discrete choice experiment survey conducted in India. The study states included Tamil Nadu (n = 98) and Andhra Pradesh (n = 83) to identify motivations driving fisher bycatch retention behaviour. Motivations to retain mobulid bycatch include: ‘Additional revenue’, ‘Bycatch preference’, ‘Reliable income’, ‘Effort’, ‘Social pressure’, ‘Food’, ‘Resource gain’, and ‘Storage space’ (Table 1).

**(a) Tamil Nadu**
| **Motivation to retain mobulid bycatch** | **Rank** | **Count score** | **Mean score** |
| --- | --- | --- | --- |
| **Additional revenue** | 1 | 255 | 2.60 |
| **Bycatch preference** | 2 | 155 | 1.58 |
| **Reliable income** | 3 | 90 | 0.91 |
| **Effort** | 4 | 6 | 0.06 |
| **Social pressure** | 5 | -75 | -0.76 |
| **Resource gain** | 6 | -201 | -2.05 |
| **Food** | 7 | -230 | -2.34 |

**(b) Andhra Pradesh**
| **Motivation to retain mobulid bycatch** | **Rank** | **Count score** | **Mean score** |
| --- | --- | --- | --- |
| **Additional Revenue** | 1 | 241 | 2.90 |
| **Storage space** | 2 | 150 | 1.80 |
| **Food** | 3 | 9 | 0.10 |
| **Social pressure** | 4 | -27 | -0.32 |
| **Bycatch preference** | 5 | -37 | -0.44 |
| **Resource gain** | 6 | -104 | -1.25 |
| **Effort** | 7 | -232 | -2.79 |

**Table 4.** Estimates from conditional logit model (best responses only) explaining fisher motivations for mobulid bycatch retention in Tamil Nadu (n = 98) and Andhra Pradesh (n = 83) in India. Fisher motivation option ‘Additional revenue’ is taken as the base case for the model. Motivations to retain mobulid bycatch include: ‘Additional revenue’, ‘Bycatch preference’, ‘Reliable income’, ‘Effort’, ‘Social pressure’, ‘Food’, ‘Resource gain’, and ‘Storage space’ (Table 1). Note: Significance indicated by asterisks p < 0.001 (***).

**a) Tamil Nadu**
|  | Estimate | Standard error | p-value |
| --- | --- | --- | --- |
| **Additional revenue** | 0.00 | - |  |
| **Bycatch preference** | -0.35 | 0.09 | < 0.001 *** |
| **Reliable income** | -0.41 | 0.09 | < 0.001 *** |
| **Effort** | -2.03 | 0.18 | < 0.001 *** |
| **Social pressure** | -2.12 | 0.19 | < 0.001 *** |
| **Food** | -3.76 | 0.41 | < 0.001 *** |
| **Resource gain** | -4.17 | 0.50 | < 0.001 *** |
| **Model statistics** |  |  |  |
| Number of respondents | 98 |  |  |
| Log likelihood | 712.8 |  |  |

**b) Andhra Pradesh**
|  | Estimate | Standard error | p-value |
| --- | --- | --- | --- |
| **Additional revenue** | 0.00 | - | - |
| **Storage space** | -0.44 | 0.10 | < 0.001 *** |
| **Food** | -0.92 | 0.12 | < 0.001 *** |
| **Bycatch preference** | -1.77 | 0.16 | < 0.001 *** |
| **Resource gain** | -2.18 | 0.20 | < 0.001 *** |
| **Social pressure** | -2.44 | 0.22 | < 0.001 *** |
| **Effort** | -5.48 | 1.00 | < 0.001 *** |
| **Model statistics** |  |  |  |
| Number of respondents | 83 |  |  |
| Log likelihood | 547.1 |  |  |

Fishers in both states chose the financial motivation ‘Additional revenue’ significantly more (p < 0.001) than all other motivators to retain mobulid bycatch (Table 4; Table 1). The top three motivators in Tamil Nadu were all financially focused: ‘Additional revenue’, ‘Bycatch preference’, and ‘Reliable income’ (Table 1). The option ‘Additional revenue’ describes the direct financial benefit, the option ‘Bycatch preference’ highlights the comparative advantage of mobulid bycatch retention than other species, and the option ‘Reliable income’ represents the value of financial security (Table 2). Fishers in Tamil Nadu chose ‘Resource gain’ as the least motivating reason for retaining mobulid bycatch. The odds of selection (calculated as the exponential of the estimate in Table 4) for ‘Resource gain’ were 98.5% lower than the ‘Additional revenue’ motivator for mobulid bycatch retention (Table 4). In contrast, the top three motivators in Andhra Pradesh were diverse, including financial and other motivators: ‘Additional revenue’, ‘Storage’, and ‘Food’ (Table 4). In Andhra Pradesh, the option ‘Effort’ was the least preferred motivator for mobulid bycatch retention, showing 99.6% lower odds of selection than the ‘Additional revenue’ motivator (Table 4).

We report significant impacts of socio-economic characteristics on fisher choice behaviour for mobulid bycatch retention (Table 5). In Tamil Nadu, fishers from Threspuram Fishing Village had a significantly lower probability of choosing the option ‘Effort’ (Table 1) than fishers from Tharuvaikulam Fishing Harbour and Tuticorin Fishing Harbour. Older fishers in Tamil Nadu were half as likely to retain mobulid bycatch due to ‘Social pressure’ (Table 1). In contrast, the likelihood of selection of the option ‘Social pressure’ increased with fishing experience. These contrasting results show a degree of heterogeneity in the responses for the option ‘Social pressure’ (Table 5). In Andhra Pradesh, fishers from Visakhapatnam Fishing Harbour were significantly less likely to choose the option ‘Bycatch preference’, than in Pudimadaka Fishing Village and Kakinada Fishing Harbour (Table 5). Similar to Tamil Nadu, older fishers in Andhra Pradesh had reduced odds of selection (approximately 33%), for the option ‘Social pressure’. Fishing vessel characteristics and operational fishing decisions also impacted mobulid choice behaviour in Andhra Pradesh. For example, fishers engaging in trips further away from coast were more likely to retain mobulid bycatch motivated by the financial option of ‘Bycatch preference’ and ‘Resource gain’. Additionally, Andhra fishers with smaller boats were more likely to be motivated by the nutritional option for mobulid bycatch retention (Table 5).

**Table 5.**
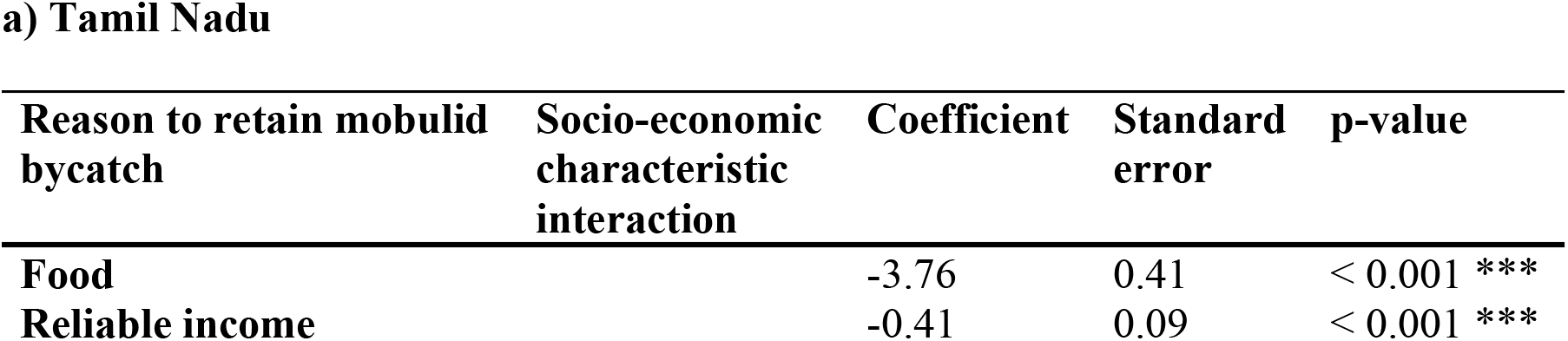
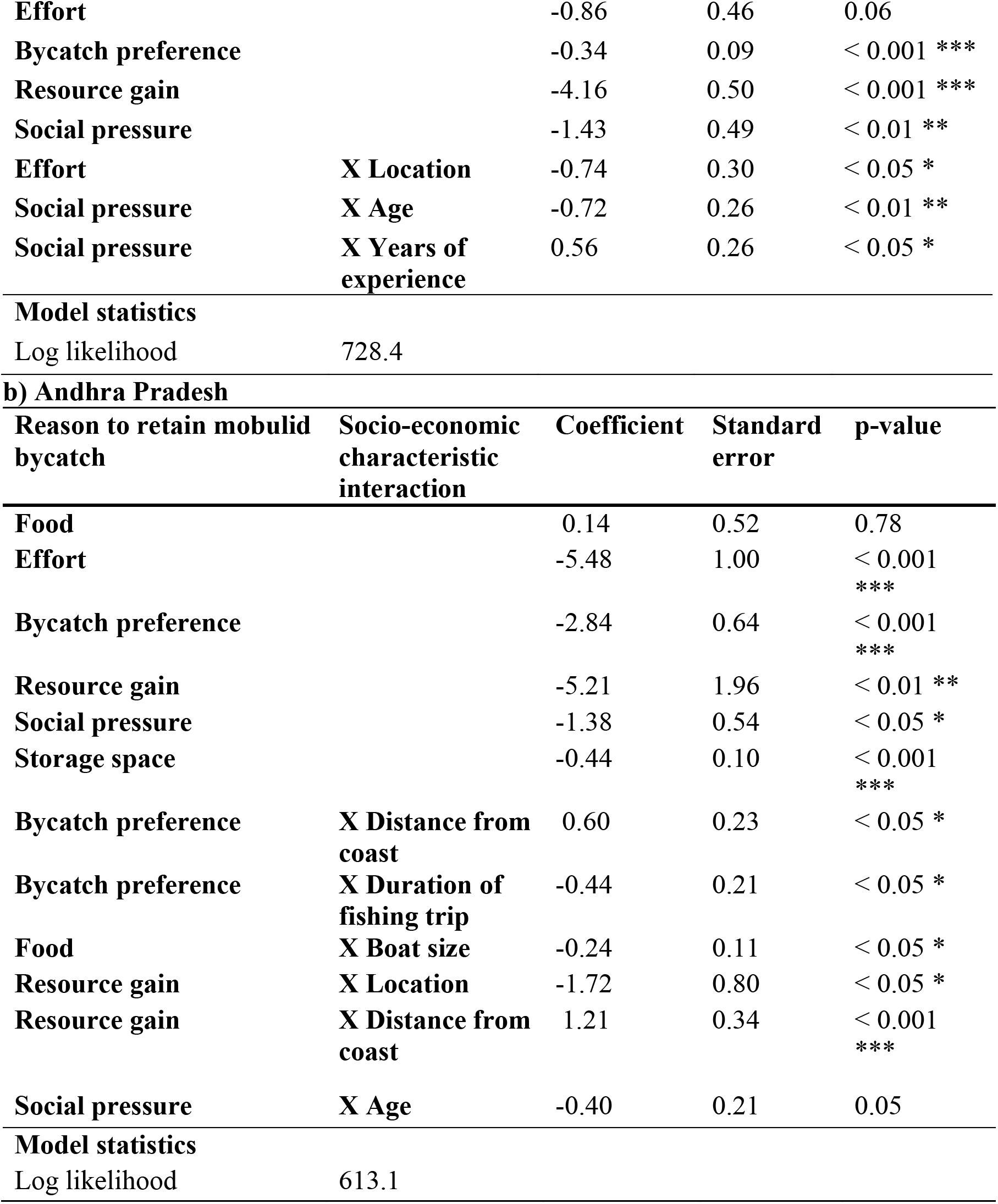
Conditional logit model (best responses only) with interaction terms describing socio-economic characteristics and reasons to retain mobulid bycatch in India. Study states include Tamil Nadu (n = 98) and Andhra Pradesh (n = 83). The interaction terms are selected with backward elimination using log-likelihood ratio values. Motivations to retain mobulid bycatch include: ‘Additional revenue’, ‘Bycatch preference’, ‘Reliable income’, ‘Effort’, ‘Social pressure’, ‘Food’, ‘Resource gain’, and ‘Storage space’ (Table 1). Note: significance indicated by asterisks p < 0.05 (*), p < 0.01 (**), p < 0.001 (***).

## 6. Discussion

Using BWS, we identified the stated motivations behind fisher choice behaviour leading to mobulid bycatch retention. We found that fishers in the fishery intensive states of Tamil Nadu and Andhra Pradesh in India exhibit varied motivations for retaining manta and devil ray bycatch in fisheries. In Tamil Nadu, the top three motivators were all financially driven: (1) selling mobulids at auction for additional revenue, (2) because mobulids are more profitable than other bycatch, so fishers prefer retaining them over other bycatch, and (3) because mobulids provide fishers with steady and reliable income because of less fluctuation in auction price. The top three motivators in Andhra Pradesh included motivations related to finances, nutrition, and storage optimisation: (1) selling mobulids at auction for additional revenue, (2) because they have space and don’t like to come back with empty storage, and (3) because mobulids are consumed as food. The financial option, ‘Additional revenue’, was consistently the highest-ranked motivation for retaining mobulid bycatch across both states. In contrast, the importance of the nutritional motivator (‘Food’) varied by state: fishers in Andhra Pradesh had approximately 60% lower odds of selecting food as a motivator relative to ‘Additional revenue’, whereas in Tamil Nadu the odds were substantially lower, with approximately 98% reduction relative to ‘Additional revenue’. Cultural differences are often responsible for spatial variation in demand, and valuation of traded products (Bachmann et al., 2019; Thomas-Walters et al., 2021). Thus, differences in key motivators for choice behaviour in our case study may be influenced by socio-cultural differences between Tamil Nadu and Andhra Pradesh fishers.

Financial dependence on fisheries for livelihoods is common in the Global South, where income from fishing often constitutes a major source of household income (Barrowclift et al., 2017; Soares and Jabado, 2024). Many coastal communities are known to derive sustenance from elasmobranch fisheries (Glaus et al., 2019; Karnad et al., 2020). Our findings show that the majority of respondents in Tamil Nadu (99%) and Andhra Pradesh (92%) are the main earners in their households. The option ‘Additional revenue’, representing direct financial benefit, had the highest relative importance for mobulid retention in both states. These results are consistent with findings from mobulid fisheries in other major catching nations in the Global South, such as Sri Lanka (Collins et al., 2023) and Bangladesh (Haque et al., 2021), as well as from other elasmobranch fisheries where communities’ dependence on elasmobranch fisheries has increased also due to overexploited stocks of other fish (Vieira and Tull, 2008; Vianna et al., 2012; Seidu et al., 2022). Additionally, bycatch utilisation patterns may be influencing the fisher bycatch retention behaviour because revenue from bycatch is often shared between boat workers, while the proceeds of the target catch may go directly to the boat owner (Salagrama, 1998). Over 70% of respondent fishers across states reported average monthly income of approximately GBP 250-270 (INR 30,000) and belong to vulnerable socio-economic groups (Umamaheswari et al., 2021), making financial dependence on the mobulid fishery even more vital to be considered in conservation policy.

Nutritional dependence of communities on shark and ray meat has emerged as one of the biggest elasmobranch fishery drivers in the past decade (Karnad et al., 2020; Pincinato et al., 2022). Our findings indicate variable importance of mobulid meat as a food source across sampled sites. In Tamil Nadu, the nutritional motivator had 98% lesser odds of selection than the direct financial motivator for mobulid bycatch retention, whereas in Andhra Pradesh, nutrition was among the top three motivators of mobulid bycatch retention. Like other elasmobranchs, mobulids are increasingly threatened by their use as a food source (Dent and Clarke, 2015). With the depletion of more traditionally targeted fish stocks, there is a growing reliance on elasmobranchs as alternative protein sources in fishing communities (Ward-Paige et al., 2012; Barbosa-Filho et al., 2019). Global meat markets further exacerbate this dependence by making access to affordable nutrition more difficult for local communities, increasing the pressure on shark and ray fisheries (Barbosa-Filho et al., 2019). However, as important as it is to recognise the stakes local communities hold in policy interventions that are equitable and context-sensitive, it is equally critical to ensure that species do not collapse or be driven to extinction (Ripple et al., 2019; Neori and Agami, 2024).

Due to socio-cultural variation, geographically nuanced incentives are essential for effective fisheries management and neglecting these differences often leads to enforcement failures and non-compliance (Spiteri and Nepalz, 2006). Our findings show geographical differences in fishers’ perceptions and preferences in our case study area. For example, the geographic variability related to the importance of nutritional benefits of elasmobranchs across states likely reflects cultural differences in dietary preferences among fisher communities (López de la Lama et al., 2018). Differences in preferences may be explained by two factors. First, there are economic disparities between respondents, with only 5% of fishers in Tamil Nadu reporting a monthly income exceeding GBP £430 (INR ∼ ₹50,000), compared to 12% in Andhra Pradesh. Second, access to international markets plays a significant role. Chennai (the capital of Tamil Nadu) is a major hub for the export of elasmobranch products (Tyabji et al., 2022; Kizhakudan et al., 2024) and is also identified as a hotspot for illegal marine wildlife trade (Lewis et al., 2022). Gill plates are potentially exported from Chennai to markets of Sri Lanka, Malaysia, Hong Kong, China, and Thailand (O’Malley et al., 2017; Kizhakudan et al., 2024). Consequently, fishery-intensive districts in Tamil Nadu, such as Thoothukudi, may have more direct access to international markets for gill plate exports. This likely contributes to the predominance of financially focused fishery perceptions and motivations for mobulid bycatch retention in Tamil Nadu.

Fisheries and trade in derived products from vulnerable elasmobranchs are causing unprecedented population declines (Hasan et al., 2023; Sherman et al., 2023), urging the need for behaviour change research in both, the product source and demand countries (Wallen and Daut, 2018; Choy et al., 2024). One of the most critical components of effective behaviour change for conservation is a clearly defined goal that aligns with the needs and values of the target demographic (Davies, 2012; Green et al., 2019; Thomas-Walters et al., 2021). Incentive-based policies have the potential to address these needs or substitute the direct or indirect values communities derive from fisheries (Leduc and Hussey, 2019; Wosnick et al., 2020). While Payment for Environmental Service (PES) schemes, such as compensation to release species (for financial motivators) are often suggested, they are neither financially viable for fisheries of this scale nor realistic as they risk increasing unsustainable behaviours (i.e., fishers targeting species during times of low-income), in addition to setting a bad precedent and crowding-out intrinsic motivations for nature conservation (Rode et al. 2015; Lee, 2022). Indirect economic incentives may include subsidies for net repair, as the release of large megafauna often results in gear damage (Bloch et al., 2016).

Our results show that in addition to direct financial motivation, fishers also value the financial security associated with mobulid bycatch retention. We recommend two potential solutions: first, the promotion of alternative species, where establishing minimum base prices for more sustainably caught fish can replicate the reliability of income that fishers seek from mobulid bycatch (Sathiadhas and Narayanakumar, 1994); second, an overall improvement in the quality of target species (e.g., through better handling and storage infrastructure), increasing their value. Additionally, awareness campaigns used to raise market desirability of seasonally sustainable species could shift bycatch retention choice behaviour to more sustainably caught species (Sun et al., 2017). Sustainable seafood initiatives (e.g., InSeason Fish) could play a key role in identifying and promoting such seasonal alternatives for substitution. Recognising the finite nature of seafood resources, we argue that sustainable seafood production must increase, or consumption must decrease, particularly among those who have the choice to do so. In parallel, broader market-based measures, such as increased awareness of the manta ray no-take policy [The Wild Life (Protection) Act, 1972 (Last Updated 1-4-2023)], the recent Convention on International Trade in Endangered Species of Wild Fauna and Flora (CITES) Appendix I listing (CITES, 2025), their listing on Appendix I and II of the Convention on the Conservation of Migratory Species of Wild Animals (CMS) prohibiting take, and the non-retention measure by the Indian Ocean Tuna Commission (IOTC), could further reduce pressure on mobulid populations.

The second critical element of effective behaviour change for conservation is the identification of a fisher population demographic that can be strategically influenced to achieve the desired positive conservation outcomes (Jones et al., 2019). Our findings indicate that fishers with more years of experience in Tamil Nadu were significantly impacted (positively) by the ‘social pressure’ to retain bycatch. This finding suggests that a campaign leveraging societal norms in Tamil Nadu would be best aimed at a more experienced fisher demographic for a higher likelihood of impact. For example, delivering the campaign through positively influential community figures could be particularly impactful (Bakti et al., 2024). For Andhra Pradesh fishers, where mobulid meat was retained more often for consumption as compared to Tamil Nadu, introducing culturally acceptable protein alternatives through campaigns led by prominent regional chefs may be particularly effective in reducing dependence on mobulids (Bowman and Stewart, 2013; Moreau and Speight, 2019).

In conclusion, identifying underlying drivers motivating fishers to retain elasmobranch bycatch is key for devising effective conservation measures. We identified that financial motivators have the most dominance for manta and devil ray bycatch retention in its largest fishery. However, the financial motivation is not limited to direct financial benefits, but is more nuanced including financial security and a comparative bycatch value with other species. As the first socio-economic study of mobulid fisheries in India, we show that motivations underlying bycatch retention decisions vary geographically and may be influenced by cultural differences between states and the socio-economic characteristics of decision makers. Based on identified fisher motivations, we provide context-specific recommendations to align conservation strategies with the values fishers derive from the fishery. We propose that identification of bycatch retention drivers can greatly aid in developing value-based incentives increasing likelihood of community adoption and thus enhancing conservation intervention effectiveness. Our approach of using BWS to identify motivations for mobulid bycatch retention choice behaviour is applicable to other mobulid priority conservation regions in the Indian Ocean, such as Myanmar, Sri Lanka, Bangladesh, and Indonesia, as identified by Palacios et al. (2025). Applying this approach broadly to other wildlife and retention behaviour contexts would enable researchers to better assess the socio-economic values derived from wildlife, further informing targeted and bycatch fisheries for improving wildlife management strategies.

## 7. Acknowledgements

The authors thank the field assistants, A. Nirojan and J. Dyson, for their invaluable help with data collection and are especially grateful to the fisher participants from the fishing villages of Tamil Nadu and Andhra Pradesh. We would also like to thank the shark and ray experts, Dr Trisha Gupta, Dr Alifa Bintha Haque, and Dr Hollie Booth, who provided feedback on the survey used in this study. We thank Rebecca Carter, Jasmine Corbett, Eithne Tynan, and Leila Scheltema from the Manta Trust team for helping to realise this work in 2024. M.C. received funding through the Swami Vivekananda Scholarship for Academic Excellence (formerly Rajiv Gandhi Scholarship), granted by the State Government of Rajasthan, Government of India. Fieldwork and data collection were funded by a 2023 Graduate Student Research Award from the Society for Conservation Biology and the Emergency Grant provided by the Manta Trust. M.C. and K.D. also acknowledge financial support from the UK Government’s Official Development Assistance by the Organisation for Economic Co-operation and Development (OECD) for the 2023–24 period.

## Notes

### Competing Interest Statement

The authors have declared no competing interest.

