## Supplementary Information for "Why are fishers retaining manta and devil ray bycatch?"

### Supplementary Information 1 - Survey development

We developed a list of 11 potential motivations for retaining shark and ray bycatch in small-scale fisheries, by conducting a thorough scoping of literature. Supplementary Table 1.1 shows the supporting references for the list of options. To reduce cognitive burden for respondents, we engaged seven experts from the field of shark and ray conservation to rank the list of options, following which the two lowest ranking motivations were removed. The remaining nine motivations were tested in pilot surveys (5 participants) and focus group discussions (5-7 participants) across both Tamil Nadu and Andhra Pradesh during the survey period in 2023 and 2024. Subsequently, based on the pilot surveys with community members, we further removed two motivations that were considered irrelevant and tested the addition of options if considered relevant by pilot survey-participants.

**Supplementary Table 1.1** Attributes and supporting references from existing literature on elasmobranch fisheries and socio-economic aspects of conservation. These were used to design the best-worst scaling survey in Tamil Nadu and Andhra Pradesh, India.

| Attributes | Supporting references |
|---|---|
| Additional revenue: Selling mobula rays at auction provides us with additional revenue. | (Glaus et al., 2019; Kizhakudan et al., 2024; Gupta et al., 2025) |
| Food: We consume mobula ray meat sometimes. | (Dulvy et al., 2017; López de la Lama et al., 2018; Glaus et al., 2019; Karnad et al., 2022; Karnad et al., 2024) |
| Cultural: We like the taste of mobula meat and cook it on special occasions and for celebrations. | (Jaini et al., 2018; Barbosa-Filho et al., 2019) |
| Effort: It saves time and effort. Removing them from the net after entanglement is difficult. | (Komoroske and Lewison, 2015; Poisson et al., 2016; Cronin et al., 2023) |
| Bycatch preference: Mobula rays are more profitable than other bycatch, so we have a preference to retain them over other bycatch. | (O'Malley et al., 2017; Muktha et al., 2024; Palacios et al., 2025) |

| | |
|---|---|
| Resource gain: The discarded parts of mobula rays are used as bait or/and poultry feed. | (Marshall and Davis, 1946; Bahadur, 2011) |
| Social pressure: I have no say in it; the entire crew works together. I don't oppose it because I will be the odd one out. | (Hatcher et al.; 2000, Alló and Loureiro, 2017; Guckian et al., 2018) |
| Storage space: They are kept because we have space and don't like to come back with empty storage. | (Carvalho et al., 2020; Roberson and Wilcox, 2025) |
| Conflict: If we don't catch them, fishers from other states and countries will catch them anyway from our waters. | (Scholtens, 2016; Manoharan and Deshpande, 2018; Amaralal et al., 2021) |
| Peer pressure: Catching big fish like mobulids is a sign of achievement in my peer group. | (Adkins, 2010; Turgo, 2014) |
| Tradition: Catching mobula rays is a tradition in our village. | (Fernando et al., 2017; Jaini et al., 2018) |

17

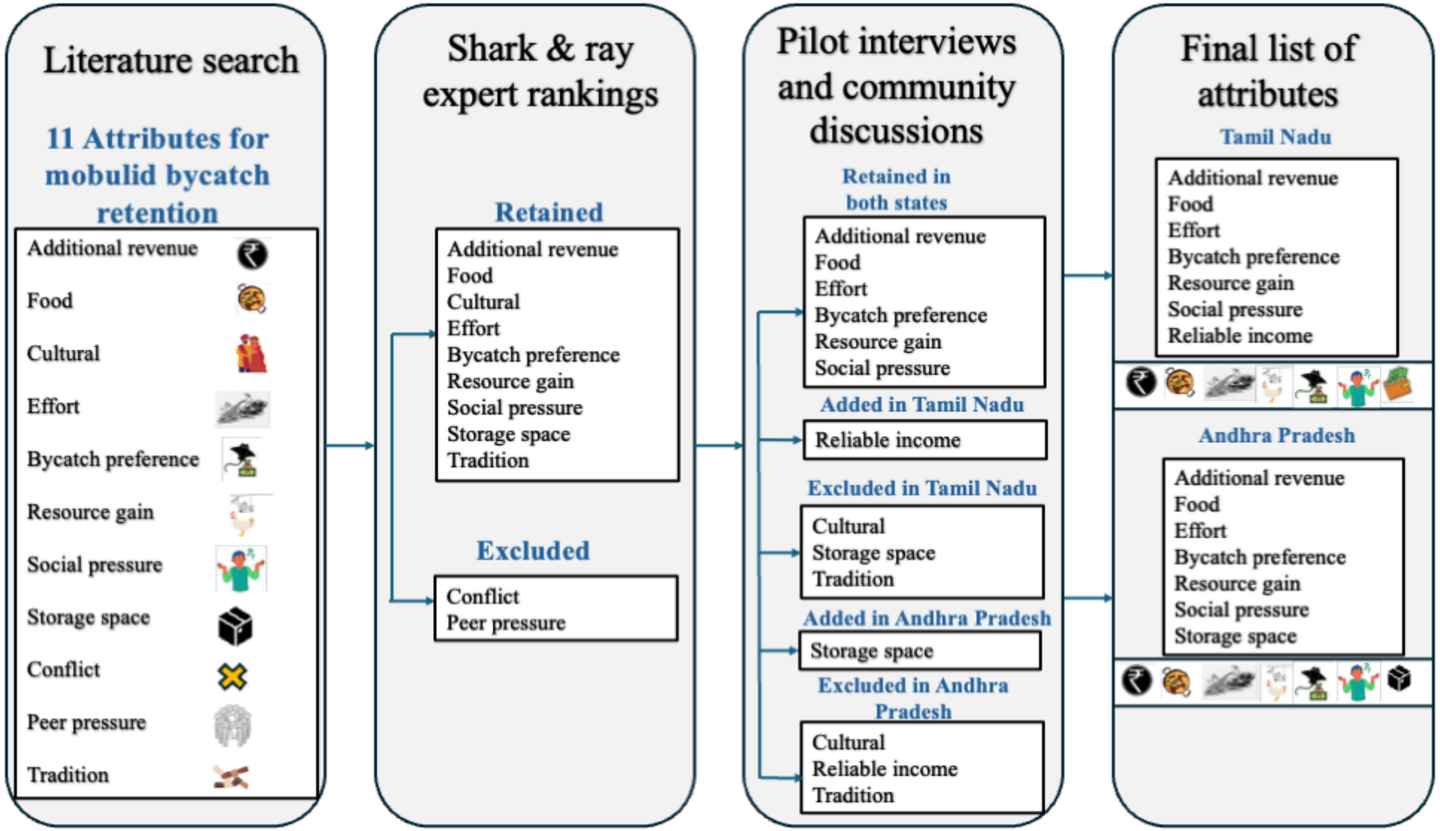


**Supplementary Figure 1.1** Survey development and attribute exclusion to develop the final list of options used in fisher surveys conducted in Tamil Nadu and Andhra Pradesh to identify fisher motivations for manta and devil ray bycatch retention.

To design the pilot surveys for the best-worst scaling experiment, we chose a 9-choice Balanced Incomplete Block Design (BIBD), comprising 12 questions with 3 options per question. Each option appeared four times across the survey and once with every other option (Louviere et al., 2015). We embedded the BIBD as 12 comparison sets in the pilot surveys. We translated the survey into regional languages (Tamil & Telugu) and had it proofread by a native speaker. The pilot surveys tested the design practicality, relevance of options, clarity of wording, and comprehension. Based on the pilot survey feedback in each state, we finalised the list of motivations. To reduce respondent fatigue, we also reduced the number of choice sets from 12 to 7 using a BIBD by Louviere et al. (2015), as participants typically lost interest after the 8th choice set.

### Supplementary Information 2 - Best-worst scaling analysis

The best-worst scores can be categorised at individual level or disaggregated scores and total level or aggregate scores (Finn and Louviere, 1992; Lee et al., 2007; Cohen, 2009). We calculated the disaggregated scores including a best-minus-worst (BW) score and its standardised score using:

$$BW_{in} = B_{in} - W_{in} \quad [1]$$

$$Standardised\ BW_{in} = BW_{in} \ / \ N_r \quad [2]$$

where $r$ is the frequency of occurrence of an object $i$, across all questions. Highly preferred options in the surveyed population result in a larger score. We calculated the aggregated score by summing up the individual-level scores across all respondents. For aggregate level scores, we defined $B_i$ was as the frequency of item $i$ being selected as the best choice across all questions for $N$ respondents; and similarly, $W_i$ as the frequency for the worst choice. $BW_i$ (which is the aggregate version of $BW_{in}$) can be given by:

$$BW_i = B_i - W_i \quad [3]$$

$$Standardised\ BW_i = BW_i \ / \ Nr \quad [4]$$

$$Sqrt\ BW_i = \sqrt{B_i\ /\ W_i} \quad [5]$$

$$Standardised\ sqrt\ BW_i = sqrt\ BW_i /\ maximum\ sqrt\ BW_i \quad [6]$$

**A. Tamil Nadu**

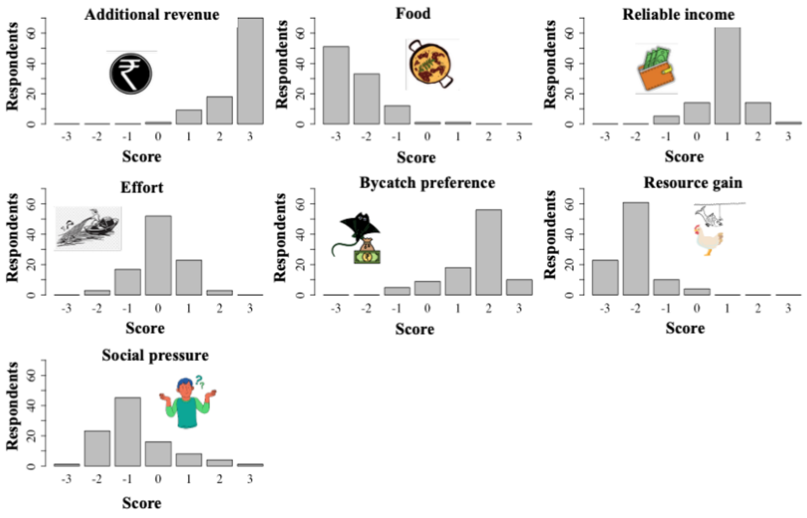


**B. Andhra Pradesh**

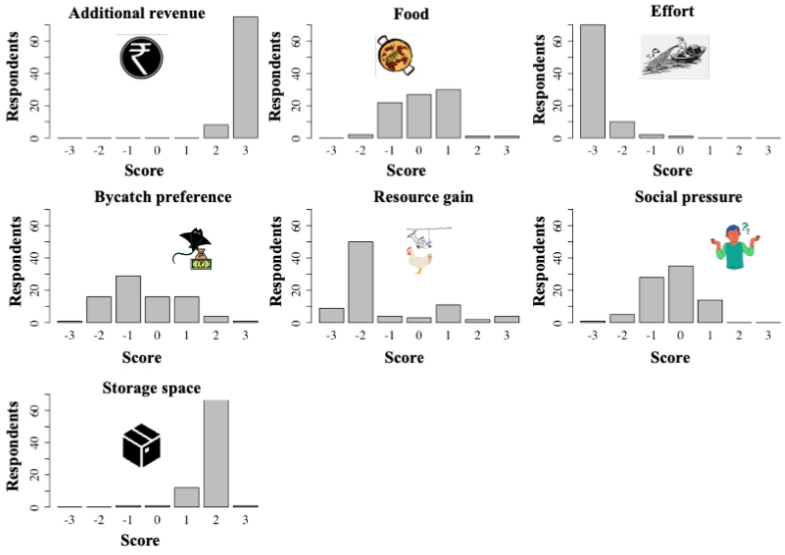


**Supplementary Figure 2.1** Distributions of best-minus-worst scores using non- parametric analysis for motivations behind fishers retaining mobulid bycatch in **A.** Tamil Nadu (n = 98) and **B.** Andhra Pradesh (n = 83), India.
